# Environmental pH signals the release of monosaccharides from cell wall in coral symbiotic alga

**DOI:** 10.1101/2022.07.03.498615

**Authors:** Yuu Ishii, Hironori Ishii, Takeshi Kuroha, Ryusuke Yokoyama, Ryusaku Deguchi, Kazuhiko Nishitani, Jun Minagawa, Masakado Kawata, Shunichi Takahashi, Shinichiro Maruyama

**Author notes:** Division of Crop Genome Editing, Institute of Agrobiological Sciences, National Agriculture and Food Research Organization (NARO), Tsukuba, Ibaraki, Japan. Corresponding author: Shinichiro Maruyama, **Email:**.

## Abstract

Reef-building corals thrive in oligotrophic environments due to their possession of endosymbiotic algae. Confined to the low pH interior of the symbiosome within the cell, the algal symbiont provides the coral host with photosynthetically fixed carbon. However, it remains unknown how carbon is released from the algal symbiont for uptake by the host. Here we show, using cultured symbiotic dinoflagellate, *Breviolum* sp., that decreases in pH directly accelerates the release of monosaccharides, i.e., glucose and galactose, into the ambient environment. Under low pH conditions, the cell surface structures were deformed and genes related to cellulase were significantly upregulated in *Breviolum*. Importantly, the release of monosaccharides was suppressed by the cellulase inhibitor, glucopyranoside, linking the release of carbon to degradation of the agal cell wall. Our results suggest that the low pH signals the cellulase-mediated release of monosaccharides from the algal cell wall as an environmental response in coral reef ecosystems.

## Introduction

Coral reef ecosystems are sustained by symbiosis between stony corals and marine dinoflagellates from the family Symbiodiniaceae, which are found in nature as free-living mixotrophs (Decelle et al., 2018; Jeong et al., 2012), as well as are primary producers in symbiotic relationships with various partners, including multicellular (e.g. Cnidaria, Mollusca, Porifera) and unicellular organisms (Foraminifera, Ciliates) (LaJeunesse et al., 2018). In oligotrophic oceans, transfer of atmospheric carbon photosynthetically fixed by the symbiotic algae to their hosts is a fundamental flux to sustain the growth and productivity of coral reef ecosystems.

Although it is generally accepted that Symbiodiniaceae algae provide photosynthates to their symbiotic partners, the molecular details are largely unknown (Falkowski et al., 1984; Ishii et al., 2019; Ishikura et al., 1999; Muscatine, 1990; Rahav et al., 1989; Stat et al., 2008; Whitehead and Douglas, 2003). Members of this family reside in an intracellular organelle called the ‘symbiosome’ within cnidarian host cells or in the extracellular ‘symbiotic tube’ systems of giant clams. These are thought to be special low pH environments that are acidified by V-type H^+^-ATPase proton pumps (Armstrong et al., 2018; Barott et al., 2015; Davy et al., 2012). While low pH environments are stressors to algae in general, they can be beneficial when CO_2_ uptake is encouraged by the hosts’ carbon-concentrating functions, enhancing photosynthesis (Armstrong et al., 2018; Barott et al., 2015). A previous study has demonstrated a photosynthesis-dependent glucose transfer from Symbiodiniaceae to sea anemone hosts (Burriesci et al., 2012), and some sugar transporters are proposed to be involved in glucose transfer (Lehnert et al., 2014; Sproles et al., 2018). Other studies suggest that the amount of transfer is regulated by the C-N balance (Rädecker et al., 2021; Xiang et al., 2020). Nevertheless, the mechanism of algal glucose secretion is not yet characterized.

## Results

To investigate the physiological effects of low pH, a characteristic environmental factor in symbioses, on algal intrinsic properties, a Symbiodiniaceae alga *Breviolum* sp. SSB01 (hereafter, *Breviolum*) was grown in a host-independent manner and cell proliferation and photosynthetic activities were measured. By comparing the growth rate of *Breviolum* in normal culture medium (pH 7.8) and acidic medium (pH 5.5, hereafter called “low pH”), we showed that the low pH medium considerably suppressed algal growth (Figure 1A) and culturing at low pH for one day resulted in significant declines in photosynthesis activity (Figure 1B).

**Figure 1.**
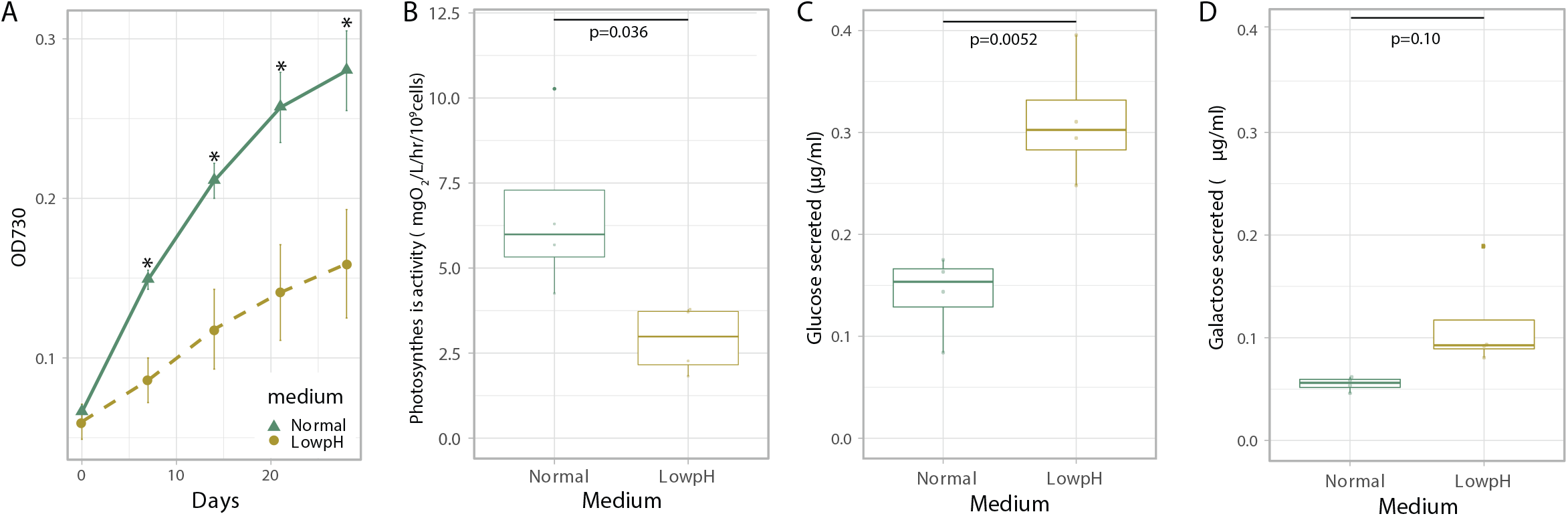
Physiological characterisation and monosaccharide secretion of cultured Breviolum. A. Growth rate (n = 6 per treatment, t-test). Asterisks indicate statistically significant differences (t-test, p < 0.005). B. Photosynthesis activity (n = 4 per treatment, t-test) C, D. Glucose (C) and galactose (D) concentrations secreted in normal and low pH media after 1 day incubation (n = 4 per treatment, t-test).

Contrary to our expectation, the amount of glucose secreted into the culture medium was higher at low pH (Figure 1C) and the secreted galactose similarly showed an increasing trend (Figure 1D). These trends suggest that *Breviolum* is capable of secreting monosaccharides autonomously without host signals, and that low pH enhanced the secretion. On the addition of the photosynthesis inhibitor 3-(3,4-dichlorophenyl)-1,1-dimethylurea (DCMU), the concentrations of glucose and galactose in the medium increased (Supplementary Figure S1), suggesting the presence of a yet uncharacterized pathway (Burriesci et al., 2012).

To investigate the response of *Breviolum* to acidic environments at the morphological level, cells cultured in different media were examined by microscopy. *Breviolum* cells in low pH media were more spread out and less clustered than the cells in normal media (Figure 2A, B). Scanning electron microscope (SEM) observations revealed that many of the *Breviolum* cells cultured at low pH exhibited wrinkled structures on their cell surfaces (Figure 2C, D). Furthermore, transmission electron microscopy (TEM) revealed that the cell surface structures of the low pH media group were ‘exfoliated’ (Figure 2E, F). These suggest that low pH affects the structures and properties of a cellulosic cell wall found in coccoid Symbiodiniaceae cells (Colley and Trench, 1983; Markell et al., 1992).

**Figure 2.**
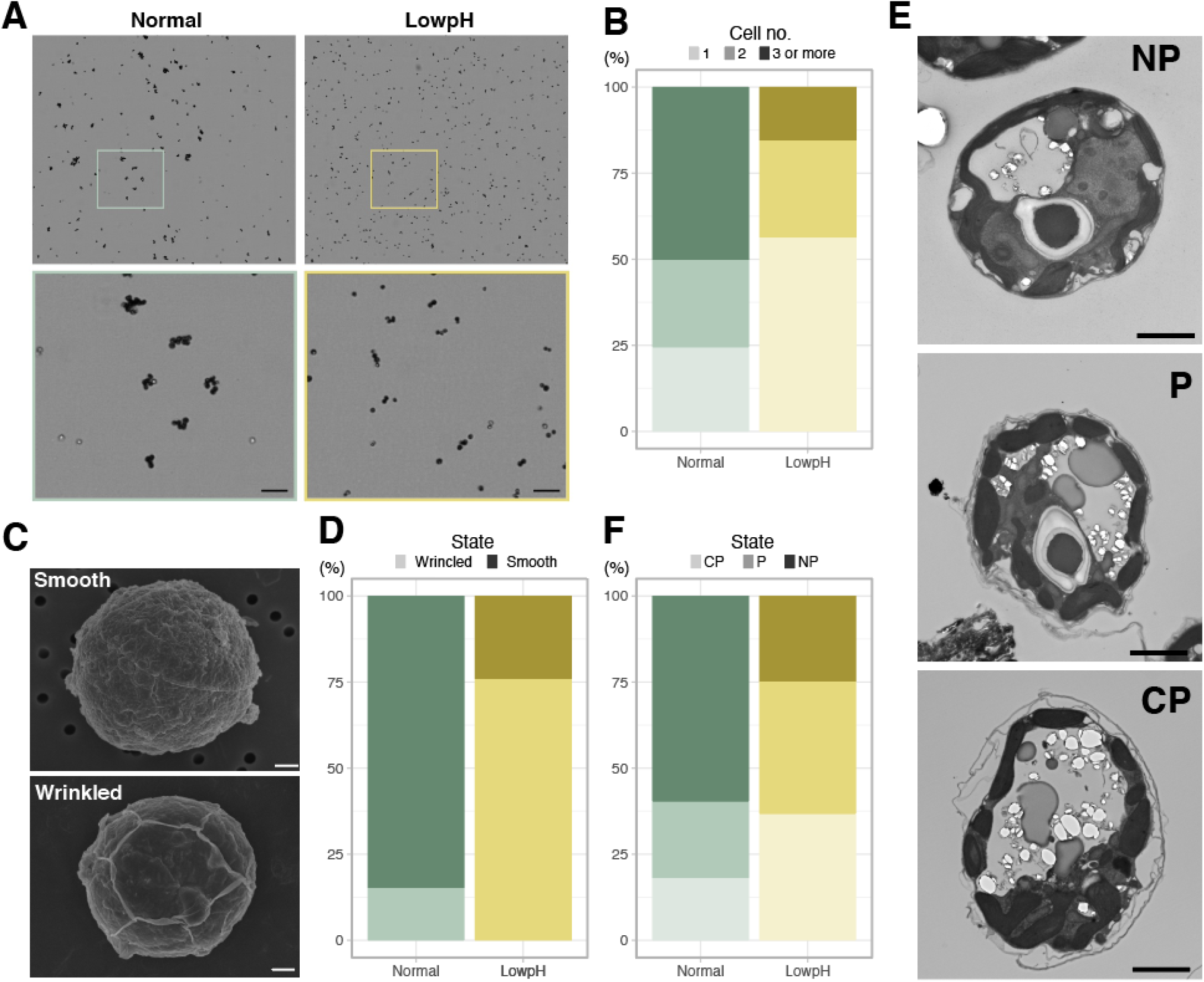
Cell structures under different pH condition. A. Bright field images of the cells under different conditions. The lower panels show high-magnification views of boxed areas in the upper panels. Scale bar = 50 μm. B. Quantification of the number of cells forming clusters (Fisher’s exact test for “1 or 2” vs “3 or more”, p = 1.727 × 10-07). C. SEM images of the representative cells. Scale bar = 1 μm. D. Quantification of the cell surface structures of the SEM images (Fisher’s exact test, p < 2.2 × 10-16). E. TEM images of the representative cells. Scale bar = 2 μm. F. Quantification of the cell surface structures of the TEM images (Fisher’s exact test for “P or CP” vs “NP”, p = 4.621 × 10-11).

To identify the mechanism involved in the monosaccharide secretion of *Breviolum*, we compared gene expression changes between the ‘control vs normal’ and ‘control vs low pH’ comparisons, and identified 3 and 4527 differentially expressed genes (DEGs), respectively (Figure 3A, Supplementary Dataset S1). The gene ontology (GO) term enrichment and KEGG pathway analysis of these two gene sets resulted in the detection of 0 (control vs. normal) and 16 (control vs. low pH) terms (Supplementary Dataset S2), which included categories related to carbon metabolism (Figure 3B, Supplementary Figure S2). The CAZy database (Lombard et al., 2014) analysis showed that 12 DEGs (28 isoforms) were annotated with Carbohydrate-Active enZymes (CAZymes) activity (Figure 3C, Supplementary Dataset S3). One of the genes, TRINITY_DN40554_c2_g2, was shown to encode cellulase (exocrine cellulolytic enzyme) harbouring a signal peptide (Figure 3C) with high similarity to dinoflagellate homologs (Kwok and Wong, 2010) (Supplementary Figure S3).

**Figure 3.**
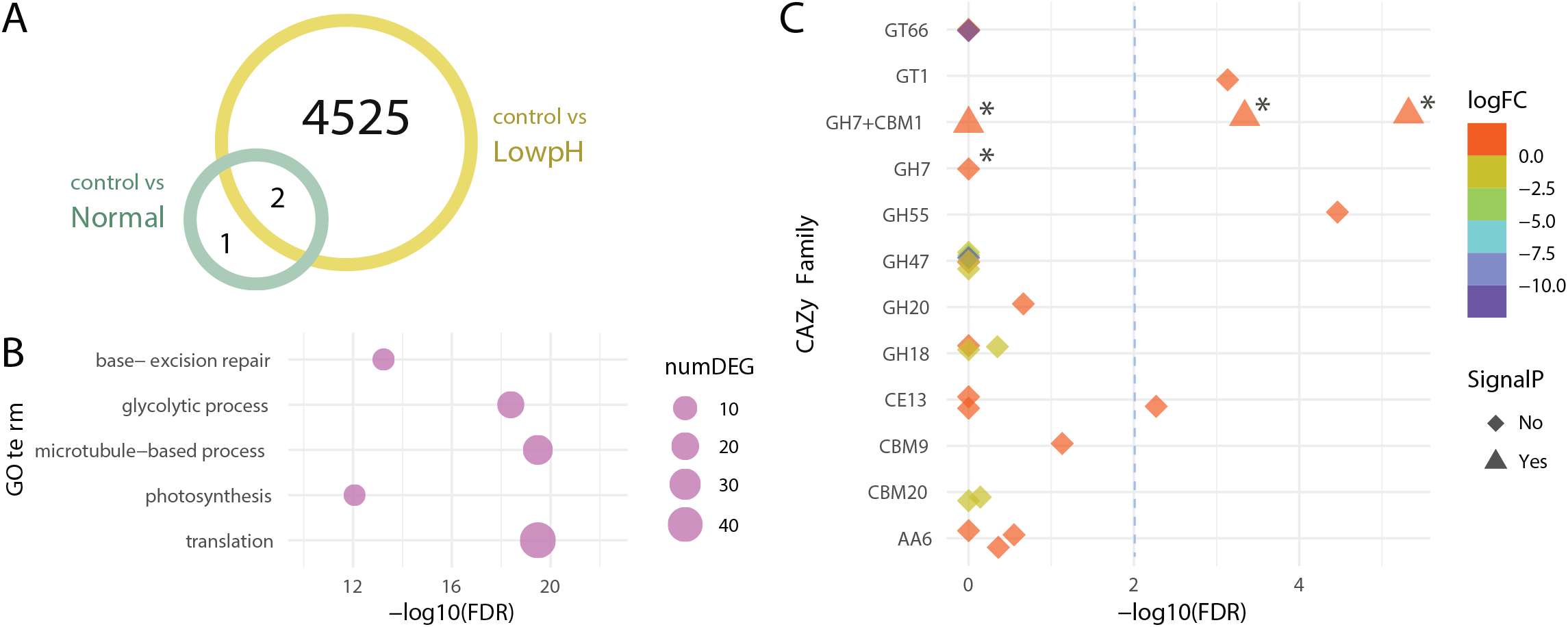
Differentially expressed genes under different pH conditions. A. Venn diagram showing the numbers of DEGs under different conditions. B. GO term enrichment analysis. Circles indicate the statistical significance (FDR) of the enriched GO terms, with the numbers of DEGs (numDEG) associated with each GO term. C. Isoform-level expression analysis of genes encoding CAZymes. Symbols indicate the statistical significance (FDR) of the differentially expressed isoforms associated with the DEGs, with the presence (triangle) or absence (rhombus) of signal peptide predicted in the amino acid sequence. Symbol colors represent the log fold-changes (logFC) of the expression levels of each isoform (low pH/control). Asterisks and dashed line indicate isoforms of the cellulase gene and a threshold for differential expression (FDR = 0.01), respectively.

To confirm the effect of cellulase on monosaccharide secretion, we examined whether secretions were inhibited by the cellulase inhibitor Para-nitrophenyl 1-thio-beta-d-glucopyranosid (PSG) (Figure 4). PSG inhibited the secretion of both glucose and galactose in a dose-dependent manner, suggesting that cell wall degradation by cellulase is involved in the secretion of monosaccharides from *Breviolum* cells.

**Figure 4.**
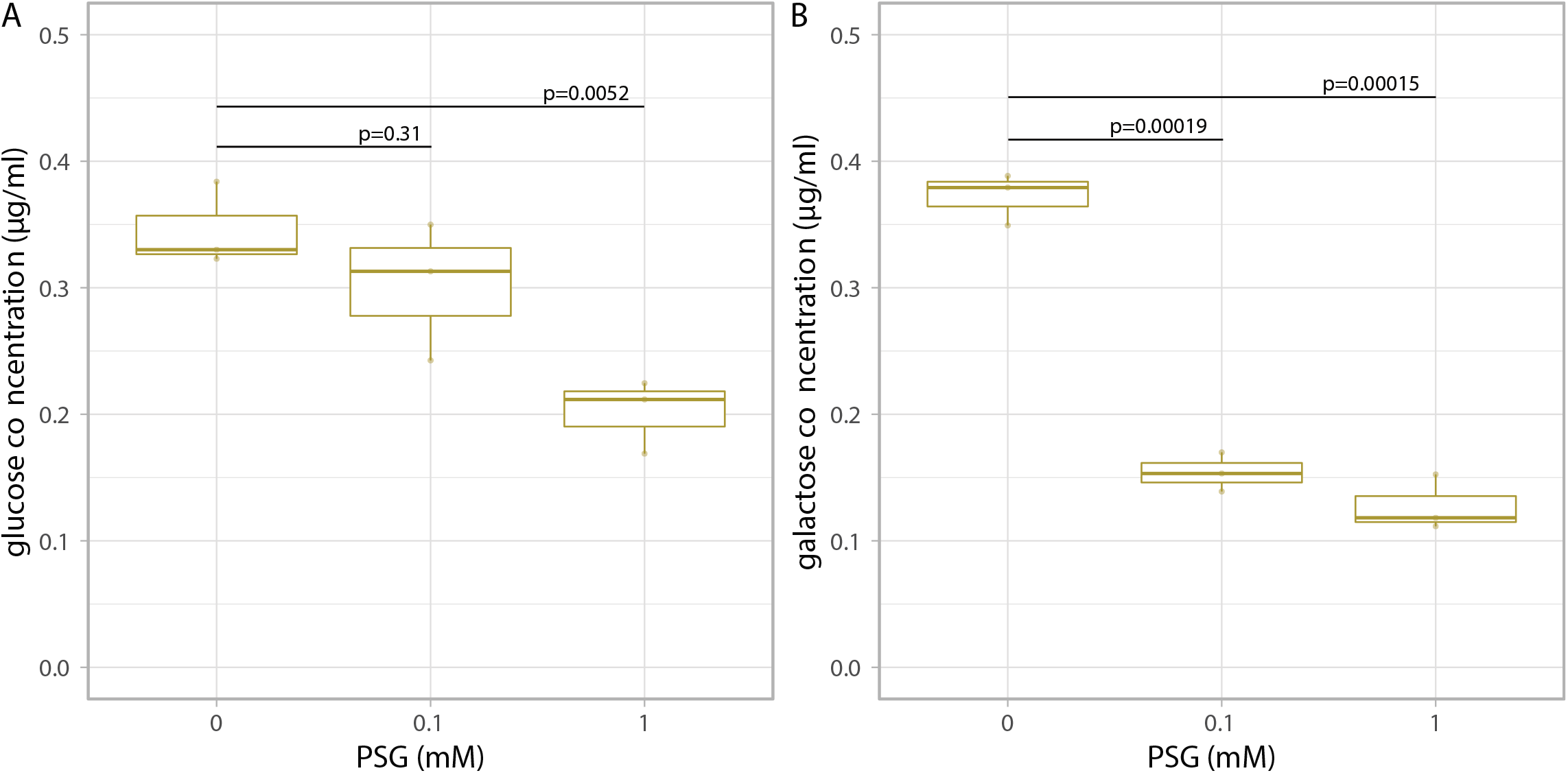
Glucose and galactose secretion in cellulase inhibitor treatment. The concentrations of glucose (A) and galactose (B) in the medium after 1 day incubation with PSG (n = 3 per treatment, t-test).

## Discussion

The transfer of photosynthetically fixed carbon from symbiotic algae to host cnidarians, including corals, is a cornerstone of their mutual symbiotic relationship (Muscatine, 1990). Unlike the current accepted model of monosaccharide secretion that assumes photosynthetically fixed carbon is directly exported from the algal symbiont via unidentified glucose transporter(s) (Lehnert et al., 2014; Sproles et al., 2018), our results suggest that stored carbon can be released from the algal cell wall as an environmental response. In the present study, we showed that a decrease in ambient pH, consistent with the interior pH of the symbiosome, accelerates the monosaccharide secretion from *Breviolum* (Figure 1). Importantly, this low pH-associated secretion was suppressed by inhibition of cellulase (Figure 4), suggesting that algal symbionts release monosaccharides into the symbiosome within host cnidarian cells by cell wall degradation. Previous studies showed that cell wall degradation/rearrangement by cellulase is required for cell cycle progression (Kwok and Wong, 2010) and cellulose synthesis is involved in morphogenesis (Chan et al., 2019) in dinoflagellates. Indeed, wrinkle and exfoliation of the algal cell wall was observed under low pH conditions using SEM and TEM, respectively (Figure 2). We need to note that our results do not deny the current accepted model, but rather suggest a multi-pathway hypothesis supported by the following observations: (i) the secretion of monosaccharides was not completely inhibited by cellulase inhibition, (ii) this new pathway occurred over days, compared to previous reports, where exported glucose was detectable in the host after only 30-60 minutes (Burriesci et al., 2012), (iii) in contrast to previous studies (Burriesci et al., 2012), DCMU did not suppress, but rather increased the secretion of monosaccharides over a longer time span (Supplementary Figure S1). Importantly, low pH upregulated the expression of genes associated with not only cellulase (Figure 3) but glycolysis probably fuelled by degradation of storage compound like starch (Supplementary Figure S2). This suggests the photosynthesis-limiting conditions trigger an environmental response in the algae to sustain cell proliferation, through cell cycle progression by cellulase, accompanying cell wall degradation and monosaccharide secretion.

Like Symbiodiniaceae, some freshwater green algae are known to be symbiotic with a range of hosts. A number of *Chlorella* strains, with and without symbiotic ability, autonomously secrete monosaccharides under low pH conditions via unknown mechanisms (Kessler et al., 1991; Mews and Smith, 1982). The monosaccharides include maltose and, to a lesser extent, glucose (Arriola et al., 2018). Dinoflagellates are also known to secrete viscous substances, including monosaccharides, as an environmental response, likely for cell aggregation and biofilm formation (Mandal et al., 2011). Thus, secretion of monosaccharides appears to be a fairly conserved environmental response and may be to compensate for retarded cell cycle progression. Therefore, acidifying symbiosomes may be an evolutionary successful strategy for cnidarian hosts to promote environmental responses of algal symbionts, which enables monosaccharides to be efficiently secreted within the organelle.

Generally, within ecosystems energy is transferred from photosynthetic primary producer to consumer by predation. Uniquely, in coral reef ecosystems energy is mainly transported from algae to corals after establishing a symbiotic relationship (Davy et al., 2012). Thus, understanding its mechanism has wider implications to understanding how energy is shared over the entire coral reef ecosystem. The multi-pathway hypothesis we propose here entails the direct transfer of photosynthates via glucose transporter(s) on their cell membrane (Lehnert et al., 2014; Sproles et al., 2018) as well as monosaccharide secretion following cell wall degradation. It remains to be determined how much each pathway contributes to the energy supply of host. However, since one pathway uses de novo photosynthates and the other uses stored photosynthates, combined they might allow for a stable supply of energy to the host, e.g., over the entire light/dark day cycle, and under photosynthesis-limiting conditions like environmental stress or cloudy days where the cellulose-related pathway could be of substantial importance. Although genetic transformation for Symbiodiniaceae is unavailable yet (Chen et al., 2019), knock-down of the cellulase gene will bring a clue to test this. Overall, our study provides a new insight into how carbon is provided by symbiotic algae to the coral reef ecosystem.

## Materials and Methods

### Strains and culture conditions

We obtained the *Breviolum* (formerly *Symbiodinium* clade B) strain SSB01, an axenic uni-algal strain closely related to the genome-sequenced strain *B. minutum* Mf1.05b (clade B), as a generous gift from Profs. John R. Pringle and Arthur R. Grossman (Shoguchi et al., 2013; Xiang et al., 2013). The *Breviolum* was maintained according to previous study (Ishii et al., 2018). Stock cultures were incubated at 25°C in medium containing 33.5 g/L of Marine Broth (MB) (Difco Laboratories, New Jersey, USA), 250 mg/L of Daigo’s IMK Medium (Nihon Pharmaceutical, Japan), and PSN (Gibco, Thermo Fisher Scientific, Massachusetts, USA), with final concentrations of penicillin, streptomycin, and neomycin at 0.01, 0.01, and 0.02 mg/mL, respectively. Light was provided at an irradiance of approximately 100 μmol photons/m^2^s in a 12 h light: 12 h dark cycle. In experiments, IMK medium containing 33.5 g/L of sea salt (Sigma-Aldrich, Merck Millipore, Germany), 250 mg/L of Daigo’s IMK Medium, and PSN with final concentrations of penicillin, streptomycin, and neomycin at 0.01, 0.01, and 0.02 mg/mL, respectively, was used. The pH of this medium was adjusted to 5.5 using HCl for the low pH experiments.

### Sample preparation for Ion chromatography, RNAseq, and electron microscopy

The incubation conditions were designed to allow detection of changes in the early phases of low pH stimulus responses. Prior to exposure to low pH conditions, *Breviolum* cultures were incubated for one week in IMK medium at densities of 8 × 10^6^ cells/20 mL in T25 culture flasks with a filter cap (TrueLine Cell Culture Flasks, TR6000) under a light-dark cycle. During experiments, *Breviolum* was cultured in two kinds of media: normal (pH 7.8) and low pH (pH 5.5), for 24 h (12 h light:12 h dark), starting at densities of 8 × 10^6^ cells/20 mL per T25 culture flask, placed at randomized positions. DCMU was dissolved in ethanol at the concentration of 20 mM and was added to cultures to a final concentration of 20 μM. The cultured cells were collected by centrifugation at 2,000 × g for 5 min at room temperature. The cells were then used for RNA-seq and electron microscopy, and the supernatants were used for ion chromatographic determination.

### Ion chromatography

Samples (n = 4 biological replicates) of the supernatant from the control (0 h), normal (24 h) and low pH (24 h) cultures were filtered using a 0.22 μm PVDF filter (Merck Millipore, Germany). These samples were loaded onto an OnGuard column (Dionex OnGuard II Cartridge) (Thermo Scientific) to remove the sulphate and halogen, according to the manufacturer’s instructions. The samples were quantified using high-performance anion-exchange chromatography with pulsed amperometric detection (HPAEC-PAD) using a Dionex ICS-5000 system equipped with a CarboPac PA1 column (Dionex) (Shinohara et al., 2017). The column was operated at a flow rate of 1.1 mL/min with the following phases: (1) a linear gradient of 0–100 mM NaOH from 0 to 31 min, (2) a linear gradient of 0–150 mM sodium acetate containing 100 mM NaOH from 31 to 34 min, and (3) an isocratic 150 mM sodium acetate/100 mM NaOH from 34 to 41 min. Myo-inositol (2 μg/mL) was added to each sample as an internal standard for quantification.

The concentrations of monosaccharides were calculated by comparing the peak ratios between the targets of interest and standards. The secreted amounts were calculated by subtracting the concentrations on day 0 from those of day 1.

### Growth rate, photosynthesis, and respiratory activity assay

To measure growth rate (n = 6 biological replicates), photosynthesis and respiratory activities (n = 4 biological replicates), *Breviolum* cultures were incubated in normal IMK media for one week and subsequently changed to fresh normal or low pH IMK media. Growth rate comparisons between the normal and low pH media conditions were conducted using 100 μL of media (625 cells/μL) in a 96-well plate. Cell growth was monitored by measuring the optical density at 730 nm (OD730) of the liquid cultures using a Multiskan GO microplate spectrophotometer (Thermo Fisher Scientific, Massachusetts, USA). Photosynthesis and respiratory activities were determined using cultures at densities of 1 × 10^6^ cells/mL in fresh normal and low pH media, on days 0 and 1 after changing the medium, separate from the transcriptome analysis. Respiration rates were calculated using the dark-phase oxygen consumption rates and photosynthesis rates were calculated by subtracting the respiration rates from the light-phase oxygen evolution rates. Mean estimates with standard errors were calculated from single measurements of four different cultures per medium condition.

### RNA extraction and sequencing

Four samples (n = 4 biological replicates) from each of the control (day 0), normal (day 1), or low pH (day 1) cultures were added to 500 μL of TRIZOL reagent (Thermo Fisher Scientific, Massachusetts, USA) and stored at −80 °C. The samples were ground with two sizes of glass beads (20 μL volume each of ‘≤106 μm’ and ‘425–600 μm’) (Sigma-Aldrich, Merck Millipore, Germany) using a vortex and performing 5 cycles of freezing and thawing with a −80°C freezer. RNA extraction with TRIZOL reagent and a high salt solution for precipitation (plant) (Takara Bio, Japan) was conducted according to the manufacturer’s instructions. The quality and quantity of the RNA was verified using an Agilent RNA 6000 Nano Kit on an Agilent Bioanalyzer (Agilent Technologies, California, USA) and a Nanodrop spectrophotometer (Thermo Fisher Scientific, Massachusetts, USA), respectively. Total RNA samples were subjected to library preparation using an NEB Next Ultra RNA Library Prep Kit (New England Biolabs, Ipswich, MA, USA) according to the manufacturer’s protocol (NEB #E7530). These mRNA libraries were sequenced in an Illumina NovaSeq6000 (S2 flow cell) in dual flow cell mode with 150-mer paired-end sequences (Filgen Inc., Japan). The raw read data were submitted to DDBJ/EMBL-EBI/GenBank under the BioProject accession number PRJDB12295.

### Transcriptome analysis

A total of 12 libraries were obtained, trimmed, and filtered using the trimmomatic option of the Trinity program. Paired output reads were used for analysis, and de novo assembly was performed using the Trinity program (Grabherr et al., 2011) to obtain the transcript sequences. The reads from each library were mapped onto the de novo assembly sequences and read count data, and the transcripts per million (TPM) were calculated using RSEM (Li and Dewey, 2011) with bowtie2 (Langmead and Salzberg, 2012).

In this study, RNA-seq produced 550,711,174 reads from the 12 samples (four independent culture flasks under three conditions), yielding 239,047 contigs by de novo assembly using Trinity. The total number of mapped reads of the quadruplicates in the de novo assembled transcriptome dataset were 77,160,394 reads for samples taken before medium change (labelled as ‘control’), 74,594,446 for the normal pH culture condition (labelled as ‘normal’), and 73,882,605 for the low pH culture condition (labelled as ‘lowpH’). Overall, we obtained the count values of the genes in the transcriptome dataset under the three conditions. Differential gene expression analysis was conducted using the count data as inputs for the R package TCC (Sun et al., 2013) to compare the tag count data with robust normalization strategies, with an option using edgeR (Robinson et al., 2010) to detect differential expressions implemented in TCC. To identify the DEGs, a false discovery rate (FDR, or q-value) of 0.01, was used as the cutoff.

To annotate the de novo transcript sequences, BLASTp search was performed (E-value cutoff, 10^-4^) against the GenBank *nr* database using all the transcript sequences as queries, resulting in 51,833 orthologs. Gene ontology (GO) term annotation of the de novo transcript sequences was performed using InterProScan (Jones et al., 2014) ver 5.42-78.0, resulting in 7,336 genes with GO terms. GO term enrichment analysis was performed using the GOseq (Young et al., 2010) package in R. Overrepresented p-values produced by GOseq were adjusted using the Benjamini-Hochberg correction (Benjamini and Hochberg, 1995). An adjusted p-value (q-value) of 0.05, was used to define enriched GO terms. In the KEGG pathway analysis, the ortholog protein sequence obtained via BLASTp search of the DEGs was used as a query. Additionally, KOID was added by blastKOALA (Kanehisa et al., 2016) (https://www.kegg.jp/blastkoala/) and mapped to the KEGG pathway using KEGmappar (Kanehisa and Sato, 2020) (https://www.genome.jp/kegg/mapper.html). For CAZome analysis, all isoform sequences of the DEGs were analysed using CAZy (Lombard et al., 2014) (http://www.cazy.org/).

### Cellulase inhibition experiment

*Breviolum* cells cultured in IMK for more than 1 week were transferred to 24 well plates at 4 × 10^6^ cells/ml density and incubated for 24 h (12 h light: 12 h dark) in IMK medium containing 0, 0.1, and 1 mM PSG (n = 4 biological replicates for each condition). The supernatant from each culture was collected following centrifugation at 2,000 × g for 2 min at room temperature. The medium samples were filtered using a 0.22 μm PVDF filter (Merck Millipore, Germany).

Glucose and galactose were quantified using a LC–MS/MS system in which a Shimadzu UPLC system (Shimadzu, Kyoto, Japan) was interfaced to an AB Sciex QTrap 5500 mass spectrometer equipped with an electrospray ionization source (AB SCIEX, Foster, CA, USA). A UK-Amino column (3 μm, 2.1 mm × 250 mm, Imtakt Corporation, Kyoto, Japan) was applied for analysis. Mobile phase A is 0.1% formic acid, and mobile phase B is acetonitrile. Samples (1 μl) were injected and analyzed over a gradient of: 0–0.5 min 95% buffer B (isocratic); 0.5–10 min 85% buffer B (linearly decreasing); 10–15 min 40% buffer B (linear decreasing). The column was equilibrated for 5 min before each sample injection. The flow rate was 0.3 ml/min. Under these analytical conditions, the retention times for glucose and galactose were 11.6 and 10.9 minutes, respectively. Mass spectrometric analysis employed electrospray ionization in the negative mode with multiple reaction monitoring (MRM) at the transitions of m/z 179→89 for glucose and galactose. The optimized MS parameters were as follows: ion spray voltage (−4500V), declustering potential (−90V), entrance potential (−10V), collision energy (−12V), collision exit potential (−11V), collision gas (N2 gas) and nebulizer temperature (450 °C). Raw data was analyzed using MultiQuant software (AB SCIEX, Foster, CA, USA). Concentrations of monosaccharides were calculated by comparing the peak ratios between the targets of interest and standards.

### Microscopy

To compare the cell conditions between the normal and low pH media, *Breviolum* was cultured and incubated for 3 weeks (12 h light:12 h dark) starting at densities of 1.6 × 10^7^ cells/20 mL per T25 culture flask. Cell photos were taken using a TC20 automated cell counter (Bio-Rad Laboratories, Hercules, CA), and the numbers of cells adjacent to and isolated from other cells were randomly counted (853 and 664 cells pooled form n = 3 biological replicates were scored in the normal and low pH conditions, respectively). The cells were fixed in a DMSO:KOH (1:1) solution, stained with 10 mg/mL calcofluor white solution in Milli-Q water, and observed under an inverted microscope (Leica DMI6000 B).

For SEM, cells were fixed in 2% glutaraldehyde and 2% osmium (VIII) oxide, dehydrated with ethanol, and dried using the critical point drying technique. The samples were coated with osmium plasma and observed under a JSM-7500F microscope at 5 kV. The surface patterns of the cells were manually scored and classified as ‘smooth’ or ‘wrinkled’ (73 and 129 cells pooled form n = 4 technical replicates were blindly scored under the normal and low pH conditions, respectively). For TEM observation, the samples were dehydrated with ethanol and embedded in EPON812 polymerized with epoxy resin. Sections 80–90 nm thick were cut, coated with evaporated carbon for stabilisation, and stained with uranyl acetate and lead citrate. The sections were then imaged at 100 kV using a HITACHI H-7600 transmission electron microscope. The cells were then categorized as NP (non-peeled), P (peeled) or CP (completely-peeled) (181 and 166 cells pooled form n = 10 technical replicates were blindly scored under normal and low pH conditions, respectively).

## Acknowledgments

We thank Drs. Naoki Shinohara and Fukumatsu Iwahashi for their support in ion chromatography and mass spectrometry, Ms Yuna Uchida for her assistance in maintaining the algal cultures, Dr. Sara E. Milward for her critical reading of the manuscript, Prof. Hiromu Tanimoto and Dr. Vladimiros Thoma for their help in preparing the manuscript, Drs. Kota Kera and Seiji Kojima for their contributions in the initial phase of the project, as well as Profs. John R. Pringle and Arthur R. Grossman for their generous gift of algal cultures. This work was supported by JSPS KAKENHI (Grant Numbers JP20J01658 [to Y.I.], JP17K15163, JP19H04713 and JP19K06786 [to S.M.]), NIBB Collaborative Research Program 18-321 and 19-332 (to S.M.), Institute for Fermentation, Osaka (to S.M.), Program for Creation of Interdisciplinary Research, Frontier Research Institute for Interdisciplinary Sciences, Tohoku University (to S.M.), and the Gordon & Betty Moore Foundation’s Marine Microbiology Initiative #4985 (to J.M.). Computational resources were provided by the Data Integration and Analysis Facility at the National Institute for Basic Biology and the NIG supercomputer at the ROIS National Institute of Genetics.

## List of figure supplements and source data files

Figure 1-figure supplement 1. Effect of photosynthesis inhibitor on glucose and galactose secretion.

Figure 1- source data 1. Raw data of growth rates.

Figure 1- source data 2. Raw data of photosynthesis activities.

Figure 1- source data 3. Raw data of glucose and galactose concentrations.

Figure 2- source data 1. Raw data of bright field cell counts.

Figure 2- source data 2. Raw data of SEM cell counts.

Figure 2- source data 3. Raw data of TEM cell counts.

Figure 3-figure supplement 1. Mapping of DEGs between the low pH and control groups on KEGG pathways.

Figure 3-figure supplement 2. Phylogenetic tree of the cellulase proteins.

Figure 3-source data 1. Expression levels of the annotated low pH DEGs.

Figure 3-source data 2. GO enrichment analysis results.

Figure 3-source data 3. CAZome analysis results.

Figure 3-source data 4. Multiple alignment of the cellulase proteins.

Figure 4- source data 1. Raw data of glucose and galactose concentrations with PSG.

---

**Figure 1-figure supplement 1.**
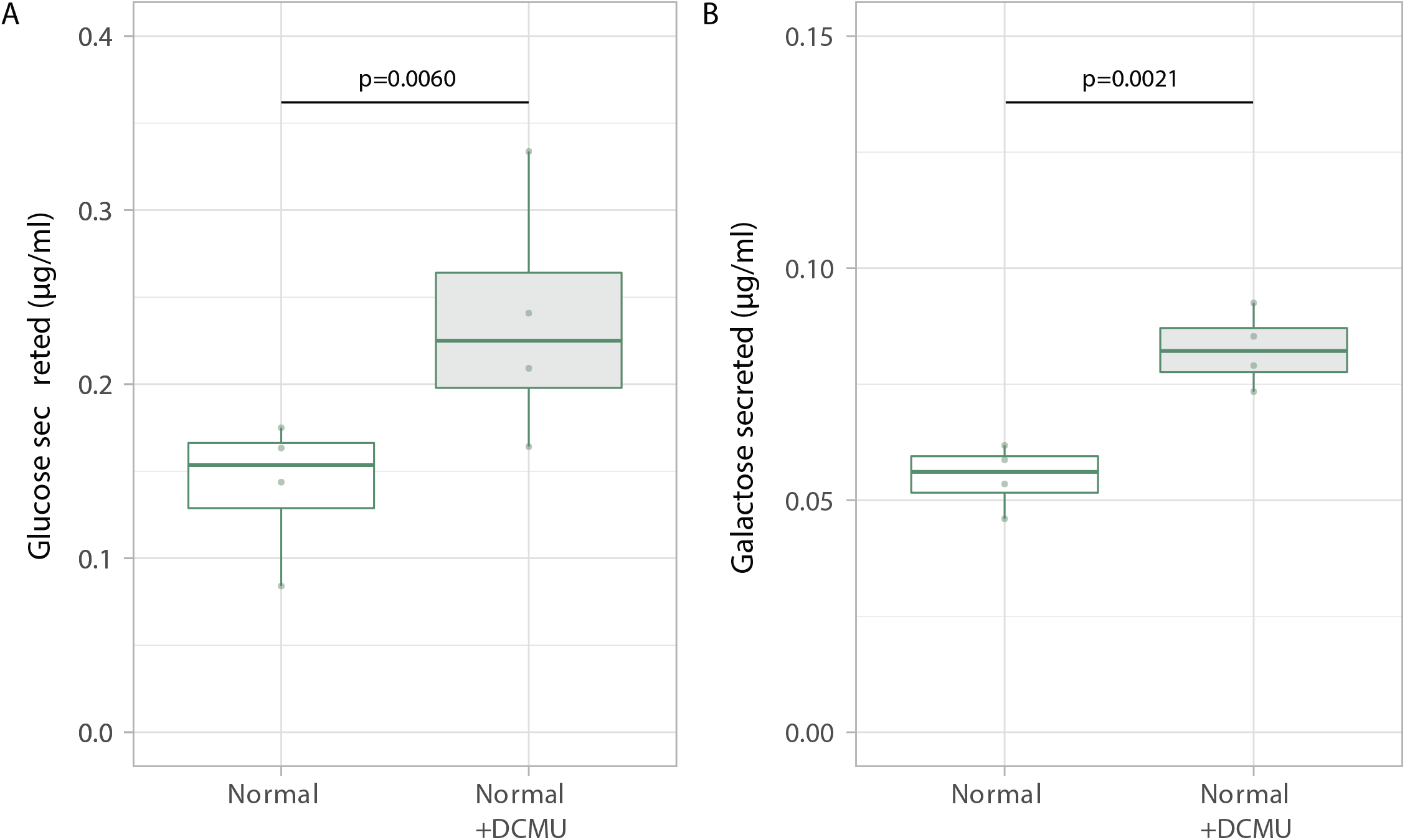
Effect of photosynthesis inhibitor on glucose and galactose secretion. Glucose (A) and galactose (B) secreted after 1 day incubation with and without the addition of DCMU (n = 4 per treatment, t-test). Values of ‘Normal’ in A were obtained from the same data shown in Figure 1.

**Figure 3-figure supplement 1.**
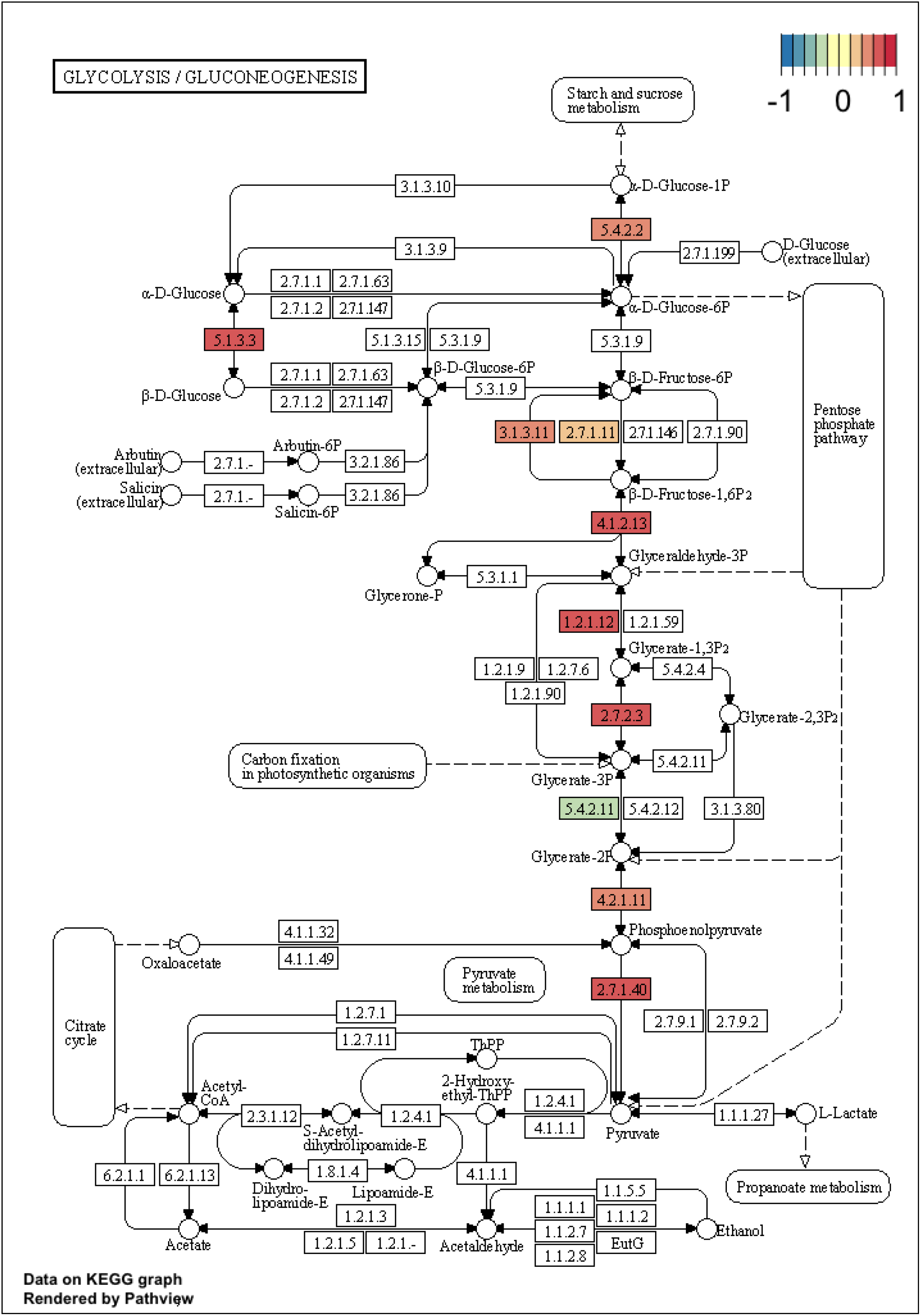
Mapping of DEGs between the low pH and control groups on KEGG pathways. The KEGG pathways for glycolysis/gluconeogenesis are shown. The colours indicate gene expression levels.

**Figure 3-figure supplement 2.**
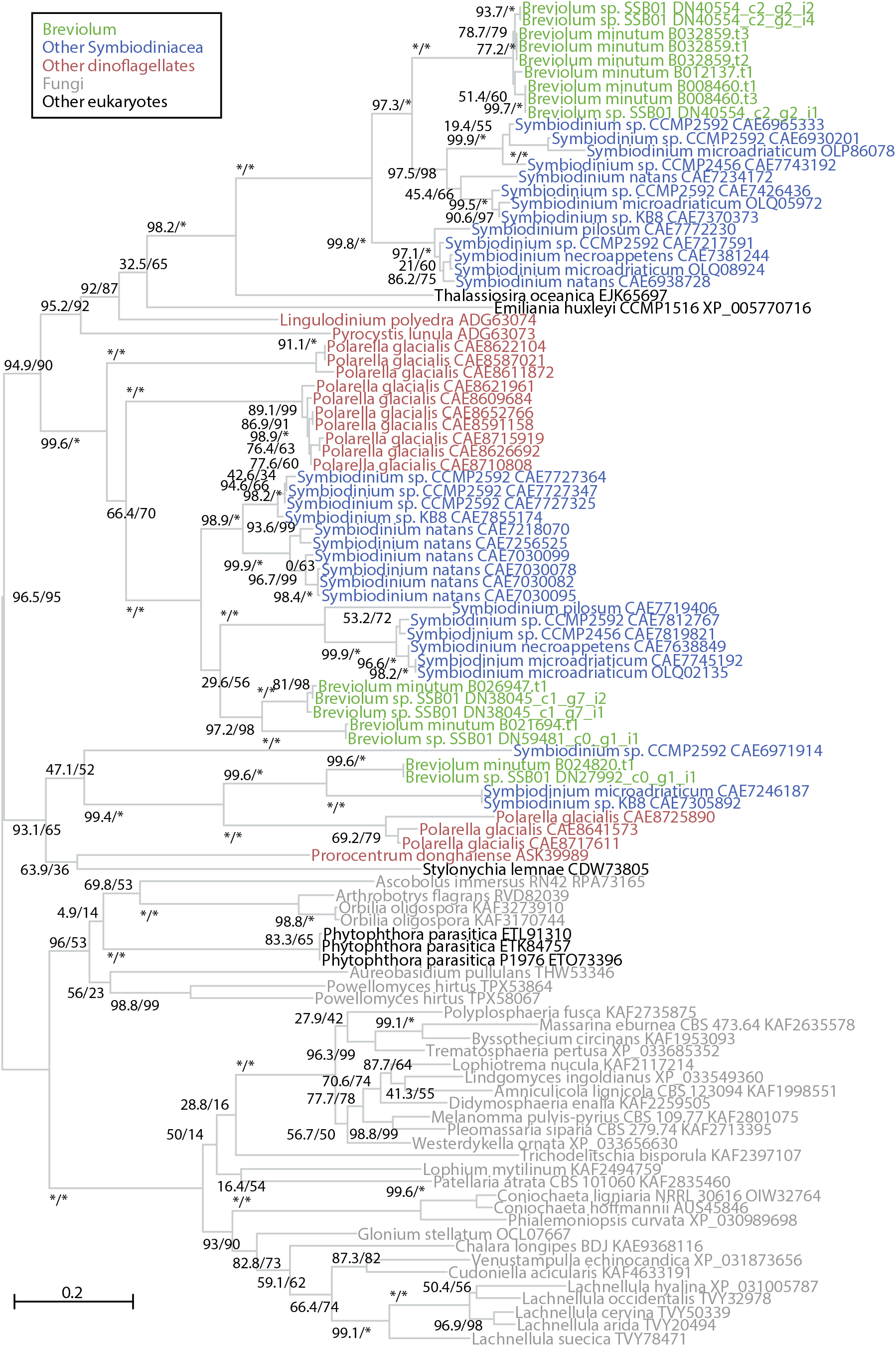
Phylogenetic tree of the cellulase proteins. A maximum likelihood tree of cellulase proteins labeled with species names and GenBank IDs is shown. Support values using an SH-like approximate likelihood ratio test (left) and ultrafast bootstrap approximation (right) are shown for each branch. For Breviolum sp. SSB01, the Trinity assembly contig IDs are shown.

